# The Bayesian-Laplacian Brain

**DOI:** 10.1101/094516

**Authors:** Semir Zeki, Oliver Y. Chén

## Abstract

We outline what we believe could be an improvement in future discussions of the brain acting as a Bayesian-Laplacian system. We do so by distinguishing between two broad classes of priors on which the brain’s inferential systems operate: in one category are biological priors (*β priors*) and in the other artifactual ones (*α priors*). We argue that *β priors*, of which colour categories and faces are good examples, are inherited or acquired very rapidly after birth, are highly or relatively resistant to change through experience, and are common to all humans. The consequence is that the probability of posteriors generated from *β priors* having universal assent and agreement is high. By contrast, α *priors*, of which man-made objects are examples, are acquired post-natally and modified at various stages throughout post-natal life; they are much more accommodating of, and hospitable to, new experiences. Consequently, posteriors generated from them are less likely to find universal assent. Taken together, in addition to the more limited capacity of experiment and experience to alter the *β priors* compared to *α priors*, another cardinal distinction between the two is that the probability of posteriors generated from *β priors* having universal agreement is greater than that for *α priors*. The two categories are not, however, always totally distinct and can merge into one another to varying extents, resulting in posteriors that draw upon both categories.

---

> *When, to silent sessions devoted to brain thought,*
>
> *We summon up formulations from endeavours past,*
>
> *And sigh the lack of many a principle that we sought,*
>
> *Because those principles were, in our mind, miscast,*
>
> *Lo, for all priors should not be tied in a single Bayesian knot*
>
> *For biological and artifactual priors have separate slots*
>
> A (posterior) Bayesian adaptation from Shakespeare’s *Sonnet 30*

## I. Introduction

We outline below a general approach to the Bayesian-Laplacian (B-L) system, which distinguishes between two major types of prior information as applied to brain studies. Our hope is that it may constitute a useful contribution to efforts in neuroscience that address the extent to which the brain uses what may be called B-L inferential operations. While we refer to the more commonly used term “Bayesian” system in the rest of this article, our title acknowledges the largely un-acknowledged contribution that Pierre-Simon Laplace (1749-1827) made in his treatise entitled *Théorie analytique des probabilités* (Laplace 1820) to the formulations in Bayes’ posthumously published paper entitled, *An essay towards solving a Problem in the Doctrine of Chances* (Bayes 1763). By speaking only of the Bayesian hypothesis, one, by implication, fails to credit Laplace with the very considerable contribution that he made in establishing the generality of the hypothesis originally formulated by Thomas Bayes.

The Bayesian approach summarizes a fundamental inferential principle, in which probabilities of occurrence of events are based on *priors* which have beliefs attached to them; through experience and experimentation, these priors lead to *posteriors*, which in turn modify inference and behaviour, thus leading to further priors with new beliefs attached to them and from which yet further posteriors may be generated (see Figure 1). The Bayesian approach is instrumental in neurobiological enquiries into how the brain’s predictive system operates, by combining prior knowledge about a phenomenon and modifying it through experience. Although we locate our interest in Bayesian operations within a broad context, namely that of the brain as a knowledge-acquiring system, we restrict ourselves here largely to addressing two questions: the extent to which experiences, in both the sensory and cognitive worlds, can update previous beliefs and lead to new ones and the extent (in terms of probability) to which the experiencing individual can assume that others, having the same experience, will update their beliefs in the same direction. The central point we try to emphasize here is that the knowledge-acquiring system of the brain and the predictions that result from it differ according to the kind of knowledge being considered. We illustrate this by referring to experiences that are significantly different, from that of colour to that of beauty, our aim being to show that the distinction we make operates at both the sensory perceptual level as well as at highly cognitive and emotional levels.

Fundamental to Bayesian operations is *belief*, which is intimately linked to priors. The brain must continually update the hypotheses that it entertains about the world, in terms of future probabilities, in light of information reaching it and against its current beliefs. Our approach leads us to enquire into different categories of Bayesian priors, the beliefs that they are based on and that they give rise to, and the role that these priors and the beliefs attached to them play in shaping the brain’s inferential systems. Our discussion is not exhaustive; rather, we hope that it lays down a basic framework for an alternative approach through which to consider the operations of the brain in a Bayesian context. The major departure in our approach is a distinction between two kinds of priors, Biological (*β priors*) and Artifactual (*α priors*). This distinction leads us to propose further that the beliefs and probabilities attached to the two categories of priors must also be distinguished according to category.

The two categories are:

1. Inherited (**biological**) or ***β priors*** are the result of (brain) concepts that we are born with; they are resistant to change even with extensive experience; the probability that the posteriors which they lead to have general agreement among all humans is high.
2. Acquired (**artifactual**) or ***α priors*** depend upon concepts which are formulated and acquired postnatally; they are modifiable and are modified by experience throughout life. They are less constrained than biological concepts and the probability that their posteriors have general agreement among all humans is low, compared to *β priors*.

As we discuss below, there are conditions in which posteriors are derived from both kinds of priors; hence the two priors are completely distinct only at the extremes.

### Definition of concepts

Of necessity, the definition of concept is different for biologically inherited concepts and for acquired, artifactual, ones. The former may be defined as inherited brain operations that are applied indifferently to incoming signals to generate biological *priors*; they could equally well be referred to as brain programs or algorithms for generating priors. We refer to the prior generated from such an inherited biological operations as the ***initial prior***, from which posteriors are generated through experience and experimentation; these then act as further (non-initial) priors for generating further posteriors. We adhere, though with diffidence, to the term *concept* because there are other, more cognitive and emotional, concepts that regulate behaviour and have beliefs attached to them, such as that of “unity-in-love” (Zeki, 2009), for which the term algorithm seems unsuitable (such concepts are not discussed further in this article). The supreme example of an initial *β prior* is in colour vision, where the biologically inherited concept, program, algorithm or ratio-taking operation (see below) is applied to incoming chromatic visual signals, regardless of their wavelength-energy composition; it results in a *colour category* which constitutes the initial biological prior for colour vision (see Figure 1). “Color [category]” in the words of Edwin Land, “is always a consequence, never a cause” (Land, 1985), meaning that it is the result of a brain operation performed on incoming chromatic signals. We devote more space to colour vision for two reasons: on the one hand we are generally clearer about the kind of inherited concept or algorithm that the brain may use to generate stable priors in colour vision and, on the other, this example can be used as a baseline against which we can discuss other categories of prior to make our point clearer.

**Figure 1:**
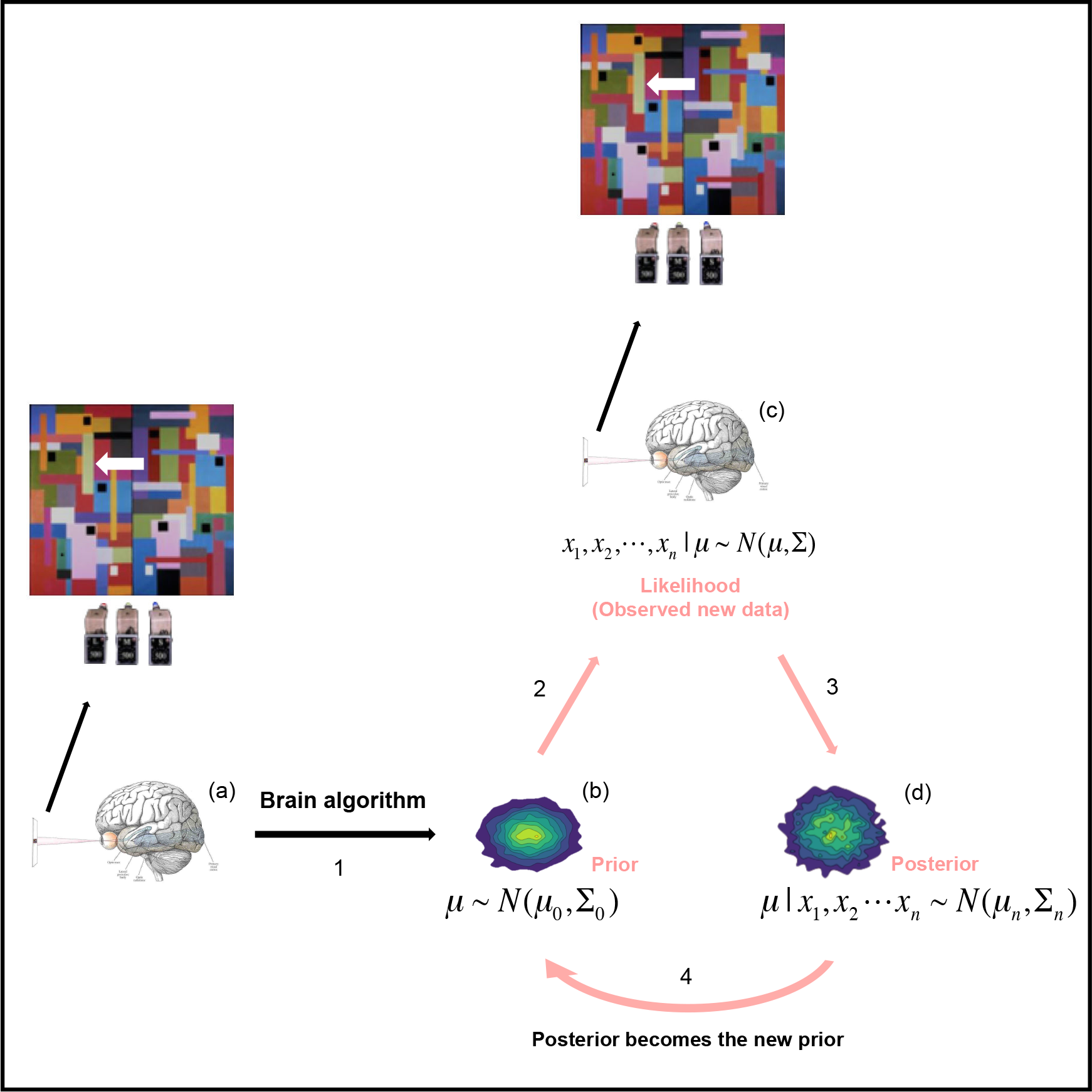
A schematic representation of the relationship of the ratio-taking process to the generation of the experience of colour. (a) It is customary to consider the brain’s ratio-taking system for colour to involve three separate processes occurring simultaneously; these are related to the ratios of the intensities of long, medium, and short wavelengths reflected from a patch being viewed and from its surrounds (the example given here is the green patch of a multi-coloured Mondrian display that is highlighted while the entire display is illuminated by three projectors, one passing long-, another middle- and the third short-wave light). This ratio-taking process generates an initial prior (as illustrated in (b)) with an initial hue attached to it. Denote the three parameters (for long, medium, and short wavelengths) as a vector μ, which follows a probability distribution (for simplicity, we assume it is a multivariate Gaussian distribution with mean and covariance μ_0_ and ∑_0_, respectively.) **(b)** Given n experiments (of viewing a patch), one observes n sets of data (x_1_, x_2_,…,x_n_), each being a three dimensional vector containing the ratios of long, medium, and short wavelengths reflected from the centre and the surrounds in each experiment i. The observed data also has a distribution (for simplicity, we assume it is a multivariate Gaussian distribution with mean and covariance μ_n_ and ∑_n_, respectively. (c) Given the prior and the observed data, the brain generates a posterior, which follows a multivariate Gaussian distribution with mean and covariance μ_n_ and ∑_n_, where 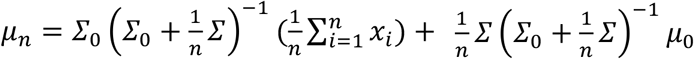 and 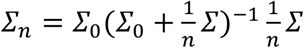. Clearly, the posterior depends both on the prior and the observed data (or likelihood in the Bayes theorem). What we argue in this article is that there exists biological priors (β priors) whose posterior is largely dependent upon the priors and independent of the observed data. Other priors fall into the category of artifactual priors (α priors), or both, where posteriors do depend upon the observed data. (d) With new experience and experimentation, the posterior becomes the updated prior, from which further posteriors can be generated (indicated by the bottom arrow).

An artifactual concept, for example that of a house or car or a game of tennis, conforms more closely to the dictionary definition of “a generic idea generalized from particular instances” (*Merriam-Webster Dictionary*). We are not born with the idea of a house or a definitive neural programme that is dedicated to the generation of houses perceptually, comparable to the ratio-taking program that generates given colour categories. Rather, the idea of a “house” is abstract and generated from viewing many houses. Moreover, the brain’s acquired concept of houses continues to grow with new experiences of houses. This is significantly different from colour vision, where the ratio-taking mechanism that leads to the generation of constant colour categories does not change with experience and learning. It is also more difficult to define what the initial prior is in the artifactual arena. This is because, unlike the inherited concept applied in colour vision, artifactual concepts are likely to change with the acquisition of experience. We argue below that certain experiences, for example that of architectural beauty, may be regulated by both acquired and inherited concepts.

Hence, in the Bayesian context, *β priors* are much more precise and intransigent to change than *α priors*, even though both can lead to almost limitless posteriors. There is good reason to suppose that the inherited, biological, *β priors* and the beliefs attached to them, which make Bayesian sense of the sensory inputs into our brains, are much more similar between humans belonging to different ethnic groups, are less yielding to experience than *α priors* and less dependent upon culture and learning than the acquired ones. Hence one cardinal distinction between the two sets of priors is that an individual can reasonably suppose that a biological *prior*, such as a colour category or the aesthetic judgment of faces or bodies as very beautiful, has universal assent, that is to say that the experiencing individual shares the judgment with the great majority of other individuals, regardless of race or culture, a characteristic not shared by the artifactual priors (*e.g*. appreciation of sushi, or a particular make of car). Specifically, the scope of experience to modify is more limited for *β priors* and the beliefs attached to them than for *α priors*. Consequently, the beliefs attached to the *β priors* and the inferences drawn from them are also more biologically constrained than the ones attached to *α priors*. In simpler terms, artifactual priors depend upon experience and experiment but biological priors do not do so or are much less dependent upon experience. Hence, we propose a modification of the classic Bayesian formula, to deal with cases where experiments are not involved in forming the prior.

## II. The need for distinguishing different categories of prior

The fundamental basis for our classification is the belief that all incoming signals into the brain are interfaced through [brain] concepts. The classification of priors into the broad categories proposed here is based in part on the Kantian system and in part upon our modification of it (Zeki, 2009). Kant wrote in *The Critique of Pure Reason* (Kant, 1781) that, “perceptions without concepts are blind”, arguing that all inputs into the mind (in our case the brain) must be somehow organized by being interfaced through concepts. In *The Critique of Judgment* (Kant, 1797) he nevertheless proposed that some sensory inputs, among them those ultimately experienced as beautiful (those that are ‘purposive without a purpose’) are either not interfaced through concepts, or only interfaced through what he somewhat vaguely termed “indeterminate” concepts; the consequence was that the perceiver usually supposed (believed), and was justified in supposing, that what one had perceived to be beautiful would also be perceived to be beautiful by others; the judgment would therefore have universal assent. This was to be distinguished from signals from objects that have a utilitarian value; the latter, Kant believed, are interfaced through “determinate” concepts and the perceiver could not assume universal assent to the qualities that one perceived in the object. There is here a logical inconsistency or even contradiction, because it is hard to see how an “indeterminate” concept can lead to an aesthetic judgment that has universal validity; surely, only a determinate concept can do so. In our system, contrary to that of Kant, it is the utilitarian, artifactual, concepts that could more easily be classified as “indeterminate” since, being dependent upon individual experience, the judgment derived therefrom cannot be assumed (in the case of the artifactual category) to have universal validity (Zeki, 2009). Thus our formulation is the reverse of Kant’s; we also differ from Kant in supposing that *all* percepts, even those pertaining to beauty, are interfaced through determinate concepts. This distinction, we believe, creates a more hospitable background for understanding the limits and capacities of how the putative Bayesian system of the brain may operate.

When considered within the framework of the brain acting as a knowledge-acquiring system, there is a further reason to emphasize a distinction between the two kinds of prior. To obtain knowledge about constant and unchanging properties of objects and situations in a world that is never the same from moment to moment, the brain must stabilize the world as best it can, a process well illustrated in colour vision, where the continual changes in the wavelength energy composition of the light reaching the brain from objects and surfaces does not, within very wide limits, modify the colour category to which they belong (see below). But this also entails having a stable prior, which codes a constant feature. If the prior belonging to the biological category were to change continually through experience and experimentation, then the capacity to stabilize the world would itself be lost because the capacity to code for a constant feature will be lost. Moreover, it is imperative for purposes of communication that the same algorithm or concept be applied in different brains to produce the same *β priors*, which can then be said to have universal assent. This also amounts to stabilizing the world, but this time doing so in terms of communication and common understanding.

The distinction that we propose is not commonly made and there is therefore no general agreement as to what disambiguates biological and artifactual priors. There are probably other attributes, besides the ones that we consider here (namely colour, faces, bodies) that fall into the biological category, for example that of landscapes and of visual motion, but we do not discuss these extensively (see, for example, (Seriès and Seitz, 2013). Rather, we give examples of what most would agree fall into different categories – colour and faces for biological categories and artifacts such as cars or planes for the acquired categories. Other beliefs, distinct from those relating to objects, also fall into the artifactual category; for example, the supposition that a government with a hardline policy on health may, if elected, lead to a rapid change in the value of companies producing medicines; these can be subsumed under artifactual priors and we do not discuss them further in any detail here since our principal interest is in priors derived from sensory inputs. Much of the distinction that we make is based on common human experience. Therefore, although the distinction between biological and artifactual priors may not have been made in the past, their phenomenology speaks to a clear qualitative distinction.

Our hope is that the differentiation we thus make will be a stimulus for further discussion on how the brain handles different categories of knowledge. We address the distinction between the proposed priors largely in terms of visual perception, about which relatively more is known and with which we are better acquainted.

## III. The Bayesian framework

The *Bayesian* framework provides analytically and biologically plausible approaches to describing how the brain operates in an uncertain world, where everything is changing from moment to moment (Gelman *et al*., 2004; Knill & Pouget, 2004a; Doya *et al*., 2007; Friston, 2012). Central to the Bayesian framework is Bayes’ theorem, namely

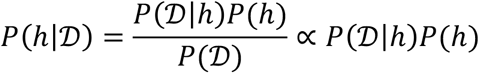

where *P* denotes probability, *h* indicates an hypothesis, and 𝒟 represents observed data. Specifically, *P*(*h*) is the prior probability of the hypothesis being true, representing our initial belief about the hypothesis before new data becomes available; thus *P*(*h*) is called the *prior probability* (or prior in short). *P*(𝒟|*h*) denotes the probability of observing data 𝒟 if the hypothesis *h* is true; it is commonly called the *likelihood function* (or likelihood in short). *P*(*h*|𝒟) represents the probability of the hypothesis being true after observing data 𝒟; it is called the *posterior probability* (or posterior in short). In the denominator, *P*(𝒟) serves as a normalization constant (that ensures the posterior distribution is a proper probability distribution) and is independent of the hypothesis *h* (and thus not useful for making a posterior inference about *h*). Thus the Bayes theorem, or *P*(*h*|𝒟) ∝ *P*(𝒟|*h*) *P*(*h*), means that the posterior depends on the observed data (through *P*(𝒟|*h*)) as well as the prior *P*(*h*). Examples using Bayesian theorem in brain studies can be found in (Tenenbaum, 1999)(Körding & Wolpert, 2004)(Murphy, 2012).

In theory, one could consider all priors under a single category which, subject to experiments or experience, will produce posteriors, as in fact previous discussions of the Bayesian brain have done (Dayan *et al*., 1995) (Rao & Ballard, 1999) (Lee & Mumford, 2003) (Kersten *et al*., 2004)(Knill & Pouget, 2004b) (Yuille & Kersten, 2006) (Friston *et al*., 2011) (Botvinick & Toussaint, 2012) (Clark, 2013) (Pouget *et al*., 2013) (Friston *et al*., 2015). The distinction we make here is, we believe, significant enough for experience to operate on the two categories in different ways. We therefore propose that the classic Bayesian formula needs to be adjusted to take account of the fact that *β priors* have very small variance between humans whereas the variance is large for *α priors*. Quantitatively, if one were to model the process within a probability framework, the variance-covariance for the prior distribution is small for *β priors* and large for *α priors*.

### III A: Colour category as a β prior that results from the application of an inherited brain concept, that of ratio-taking

For colour, the Bayesian theorem would state that there exists a ratio-taking mechanism or algorithm (Land, 1974) (Land, 1986) (Land & McCann, 1971)(Zeki, 1984) which is very largely independent of culture, learning and experience, and is therefore identical (*i.e*. with very little variance) in all humans. This algorithm generates an initial prior which is the colour category, to which an initial hue is attached. From this posteriors are generated through experimentation, for example by viewing a patch belonging to a given colour category (say, green) under different illuminants or with different surrounds. This can be illustrated by the following example: if the initial (green) rectangle in Figure 1 is viewed under a different illuminant, its hue will change, although its colour category will remain the same. Hence the colour category is relatively immune to experience and experimentation while the initial hue is much less so. Hence, too, by illuminating the patch with light of different wavelength composition or by modifying the colour of the patches that surround it, many posteriors in terms of hue can be generated from this initial prior. With each new experience and experimentation, the posterior becomes the updated *prior*, from which further *posteriors* can be generated (see Figure 1).

These considerations lead us to propose the following *Bayesian Brain Theorem* (see Supplementary Information and Figure SI) for mathematical details, which also apply to the generation of other biological priors) which summarizes the above example theoretically: the *β prior* (*e.g*. the *constant colour category* generated from the ratio-taking operation detailed above) has attached to it another *β prior*, which is the hue. Let us refer to this as 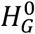, where the subscript *G* refers to the green colour category, and the superscript to its initial hue. The experiment conducted (for example, varying the wavelength-energy composition of the illuminant in a colour experiment or surrounding the green patch with a patch of a single colour, say red) will lead to a posterior hue which, though still belonging to the colour category of green, will differ in its shade of green (and hence hue) from 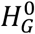. We will refer to this as 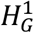. As one experiences different shades of green (different hues) when one views the same scene in different illuminants or, in the example of the green patch above, changes its surrounds, and notes the nature of the changes in the illuminant and/or the surrounds of the green patch being studied, so any number of different shades (hues) of green can be generated and experienced. These posteriors (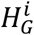, where *i* = 0, 1, 2, …,) can be updated continuously and iteratively in one’s life; knowing more about the illuminant or the spatial configuration of a stimulus, one can therefore make inferences with a high degree of accuracy and reliability (see the iterative process in Figure 1).

Colour categories represent perhaps the most extreme form of a *β prior* which is the result of an inherited brain concept applied indifferently to received chromatic signals to generate constant colour categories (see below). It is more usual, in the psychological literature, to speak of colour constancy; we prefer to speak of the ***constancy of colour categories*** (see Zeki *et al*., 2017); we do so because the precise hue (shade of colour) within a (constant) colour category varies with changes in the wavelength composition of the illuminant, but the colour category itself does not; hence it is more accurate to speak of a constant colour category (see Zeki *et al*., 2017; 2018).

To summarise, colour, or more precisely a colour category, is an experience; the initial experience generated from a ratio taking mechanism is that of a colour category to which a given hue is attached. Through different experimentations, further hues (shades of colour) may be generated; these further hues, though belonging to the same colour category, differ in appearance (in shade of colour) from the initial hue, depending upon factors such as the wavelength composition of the light reflected from, say, a green surface and the colours surrounding it (see Figure 1). These further hues thus become *posteriors* which then act as new *priors* from which yet further hues (*posteriors*) can be generated through experiments and experience (see the iterative process in Figure 1). In brief, the constant colour category (green) with an initial hue attached to it together constitute the primal experience of colour (*β prior*). While the colour category itself remains largely resistant to change through experience and hence its posteriors are little modifiable despite varying experimentations, further posteriors of the hue can depart from the initial hue attached to the colour category; the direction in which the posterior (hue) changes can be predicted with high accuracy through experience (see below); it is indeed this knowledge gained through experience and experimentation that artists use constantly. In this context, we note that the cortical response to colour in pre-linguistic infants (5-7 months) measured by near infra-red spectroscopy indicates that there is a significant increase in activity in occipito-temporal regions (presumably including area V4) with between-category (colour category) alterations but not with within-category (hue) alterations (Yang *et al*., 2016); similar results have been reported in other studies comparing infants and monkeys (Bornstein *et al*., 1976), consistent with the view expressed here, that constant colour categories are the *priors* to which the initial hue is attached, and from which further hues (posteriors) may be generated.

Constant colour categories and the hues attached to them are biological signalling mechanisms allowing the rapid identification of objects. If the colour category of an object or surface were to change with every change in the illuminant in which it is viewed, then colour would no longer be a useful biological signalling mechanism, because the object can no longer be identified by its colour alone. To make of colours an autonomous identifying mechanism, they must be stabilized and be immune, as far as possible, from the de-stabilizing effect of a change when the wavelength composition of the illuminant in which objects and surfaces are viewed changes. This fulfils one of the important requirements when considering the brain as a knowledge acquiring system, to which is coupled the need to stabilize the world. We also presume, as stated above, that the same processes operate in different brains to generate the same constant colour categories, making it easy for an individual to assume their experience of a constant colour category has universal assent, thus fortifying another reason for having *β priors* with small variance, namely easing the process of communication between individuals.

Evidence shows that there are specific brain pathways and a specific visual area, area V4 and the associated V4a, that are crucial for colour perception (Zeki, 1973)(Zeki, Watson, & Lueck, 1991)(Bartels & Zeki, 2000), damage to which leads to the syndrome of cerebral achromatopsia (Meadows, 1974)(Zeki, 1990). It is important to emphasize that the cells of V4 not only respond specifically to categories of colour but also to hues, or shades of colour (Zeki, 1980)(Stoughton & Conway, 2008) (Brouwer & Heeger, 2013) and that the representation of colour within V4 can be independent of form (Zeki, 1983)(Lafer-Sousa, Conway, Kanwisheer, 2016). It is therefore likely that area V4 is pivotal to these operations (Bartels & Zeki, 2000), which is not to say that it acts in isolation; it does so in co-operation with the areas it receives signals from and projects to, together with the reciprocal connections between these areas.

### III B. The brain’s ratio-taking system for generating constant colour categories and the constants in nature

Colour (and the category that it belongs to) is a brain construct (Zeki, 1984); it is generated through an operation based on an inherited concept (algorithm or program), that of ratio-taking, described above. Many different ways of implementing this have been proposed (for a review, see Foster 2011) but they all share a common feature, namely a comparison of the wavelength composition of light reflected from different surfaces. This is what we, too, emphasize here although we rely more on the classical approach of Land and his colleagues without implying that it is the final word on the implementation. Using the arguments presented here and the mathematical explanation given in the Supplementary Information, we show that *this process is independent of any experiment or experience and that a* colour category (as opposed to a shade of colour or a hue) is only dependent upon the ratio-taking operation of the brain (see Figure 1). It is useful to discuss briefly here why this should be so in the context of what the constants in nature are and how these constants generate constant colour categories.

The unvarying property of surfaces in terms of colour vision is their reflectance, namely the amount of light of any given waveband – in percentage terms – that a surface reflects in relation to the light incident on it. For a given surface, this percentage never changes. Hence, one finds that the ratio of light of any waveband reflected from a given surface and from its surrounding surfaces also has little variability, regardless of the variation in the intensity of light of that waveband reflected from it; if the intensity of light of any given waveband reflected is increased, the intensity of light of the same waveband coming from the viewed surface and its surrounds also increases, and the ratio continues to remain the same. By extension, the ratios of light of all wavebands reflected from a surface and from its surrounds also have little variability. For a more detailed (and quantitative) explanation see Supplementary Information.

Take as an example a green surface which forms part of multicoloured (natural) scene, as in Land’s colour Mondrians; it is surrounded by many patches of other colours (Figure 1 (a) shows a much simplified version). Let us suppose that the green patch (*g*) reflects *x* per cent of the long-wave (red) light, *l*, incident on it, *y* per cent of the middle wave (green) light, *m*, and *z* per cent of the short-wave (blue) light, *s*. The surrounds, having on average a higher efficiency for reflecting long-wave light, will always reflect more and there will be a constant ratio in the amount of red light reflected from the green surface and from its surrounds. Let us call this ratio *g*^*l*^. The surrounds will have a lower efficiency for reflecting green light and hence there will be another ratio for the amount of green light (*g*^*m*^) reflected from it and from the surrounds, and a third ratio for the amount of blue light *g*^*s* 3^. When the same natural scene is viewed in light of a different wavelength composition, the amount of light of different wavelengths reflected from a surface and from its surrounds will change, often significantly (as, for example, when a scene is viewed successively in tungsten light, in fluorescent light, or in sunlight) but the ratios in the amount of light of different wavebands reflected from the centre and from the surrounds remain the same.

Formally, let us consider the green patch viewed under two different illuminants, where the first one has 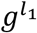, 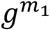 and 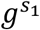 amounts of long, middle, and short wave light reflected from the green patch, and the second one has 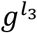, 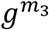 and 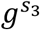 reflected from it. The first illuminant results in (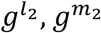, and 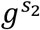) and the second in (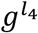, 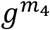 and 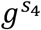) ratios of long-, middle-, and short-wave light reflected from the green patch and from its surrounds, respectively. Then, we have:

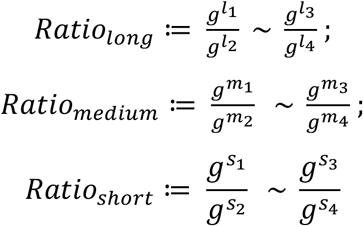

where ~ means that the long, medium, and short ratios differ little (have very small variance) from different experimental settings (namely, regardless of the precise wavelength-energy composition of the light reflected from the green patch and from its surrounds). More concretely, let µ = (*Ratio_long_*, *Ratio_medium_*, *Ratio_short_*). Denote every individual *i*’s ratio-taking mechanism at time *t* as µ_it_, which is sampled from a probabilistic distribution. We argue here that the distribution of µ_it_ has a very small variance in time and between humans. Suppose µ_it_ is generated from a multivariate Gaussian distribution with mean and covariance µ_0_ and Σ_0_, respectively. Then, that there exists a *β* prior for colour is equivalent to saying that the entries of Σ_0_ are very small (see Supplementary Information for more information). In other words, the ratio-taking mechanism varies little from individual to individual, and from time to time.

It is through such a ratio-taking mechanism that the brain builds *constant colour categories*. We have no knowledge of the detailed neural mechanisms by which the brain implements this but there is no physical law that dictates that such ratios should be taken or comparisons made; it is instead an inherited brain law and, given the widespread use of constant colour categories in the animal kingdom, we make the assumption that similar mechanisms, or very nearly so, are used in species as far apart as the goldfish (Ingle, 1985) and the human (Land, 1974). These constant colour categories constitute the *β priors*. This generation of constant colour categories is a classic example of Kant’s statement in his *Prologemena* (Kant, 1783) that “The Mind [brain] does not derive its laws (*a priori*) from nature but prescribes them to her”.

No amount of visual experience can modify the colour categories (*β priors)*; in fact they often cannot even be modified by higher cognitive knowledge (but see also below). For example, green leaves may reflect more green than red light at noon and more red light than green at dawn and at dusk; but they are always perceived as green, even in the face of knowledge that they are reflecting more red light under certain conditions, provided the leaves are being viewed in a natural context. Technically, colour categories are therefore endowed with precision. Priors exert a more constrained effect over posteriors when they are more precise, as in biological priors. We use the term precision here in a technical fashion, to denote the confidence afforded to priors; precision is the inverse dispersion or uncertainty encoded by a probability distribution (*e.g*., the variance of a Gaussian distribution). We define precision as the inverse of variance. In normal colour vision, given the *β prior*, the variance is very close to 0 and the precision therefore approaches an infinite value. Hence the *β priors* in colour are extremely stable and un-modifiable with experience. It goes without saying that (language apart), an individual experiencing a given colour category C is entitled to suppose that his or her experience is universal, in that all other (normal) individuals will experience the same colour category under the same conditions of illumination. This can be readily established by matching the colours perceived with Munsell chips, without the use of language (Zeki *et al*., 2018). Hence the belief is universal. Because of this assumed universally shared belief, Kant would no doubt have referred to this as an experience which is not interfaced through a (prior) concept or one that is interfaced through an “indeterminate” concept. But, as we have tried to show, it is in this instance an experience generated through an inherited ratio-taking operation, a determinate inherited concept operating without regard to race, culture and upbringing, and executed by the brain to stabilize the world in terms of colour categories. We leave out of account here the vexed and unsolved problem of qualia, of whether the quality of green that one person perceives is identical to that perceived by another.

We have gone into colour in some detail because it best illustrates how the brain can stabilize the world through the generation of an initial *β prior* without the need to experiment or acquire experience.

### III D. The ‘belief’ with respect to colours

A definition of ‘**belief**’ with respect to colours might be adapted from its ordinary dictionary definition, namely “a feeling of being sure that someone or something exists or that something is true” (*Webster’s Dictionary*), or “confidence in the truth or existence of something not immediately susceptible to rigorous proof “(*Dictionary.com*) or that “the experience (of colour) will always be true”, even when we are not remotely aware of the operations that lead to the experience. The belief with respect to colours is subtle. Both Hermann von Helmholtz and Ewald Hering tried to account for colour constancy by invoking higher cognitive factors; Helmholtz (1911) by invoking judgment and *a priori* knowledge, believing that the viewer assigned a constant colour through an adjustment due to the vaguely defined process of the “unconscious inference” and Hering (1877/1964) by supposing that memory was essential. While memory and learning may play a role and modify the experience of colour for objects about which one has knowledge through experience, this is, significantly, not true for colour that is detached from definitive objects (Vandenbroucke *et al*., 2016) or colour attached to “nonsense” objects, of which Land’s experiments constitute a classic example. But even such knowledge may be over-ridden by the brain’s computational process to generate colours, as Land’s two colour projection experiments show; in these, the identical picture of a multi-coloured natural scene is taken through different filters (say a long-wave and a middle-wave filter). The (black and white) pictures produced are then projected in perfect alignment on a screen, one through a projector with white light and the other through a long- or middle-wave filter. When so projected, a normal viewer will see a variety of colours. The colour of the objects with which the viewer is familiar is not determined by *a priori* knowledge but rather by the filter used, in combination with white light, to project the image; lips, for example, may appear blue even in spite of knowledge that they are not normally so (Land, 1959). Helmholtz qualified his “unconscious inference” in a way from which we depart, for he supposed that judgment and learning enter into that inference. Along with Land, we prefer to think of a computational process that is independent of such cognitive factors and that only requires a comparison of the reflectance properties of surfaces. If learning, judgment, memory and experience are relevant here, as claimed by both Helmholtz and Hering, it is only with respect to the precise hue or shade. A viewer ‘knows unconsciously’ that the green patch will look green if it reflects more green light than its surrounds, regardless of the actual amount of green light reflected from it and regardless of whether one is acquainted with the object or not, thus precluding judgment and learning. Anyone armed with such technical knowledge (though of course few are) can predict the colour of a surface, even before seeing it. Thus, a red surface is one that will reflect more red light than its surrounds and a blue surface is one that will reflect more blue light than its surrounds. A white surface will reflect more light of all wavebands than its surrounds while a black one will reflect less light of all wavebands than its surrounds, regardless of the actual amount of light reflected from it. This belief system is quite rigid and not easy to manipulate.

## IV. Faces – another category of *β* priors

We argue below that faces also belong to the category of *β priors*. But if reflectance is the constant in colour perception, the constants in face perception are more difficult to determine and define. We may say that there are certain elements such as the nose, mouth, eyes, and so on (not all of which have to be present at once) that must be present in a harmonious and proportionate relationship to each other; these constitute an essential arrangement of parts which we refer to as a “significant configuration” (Zeki, 2013) and which leads to the instant recognition of a face as a face. The elements constituting that arrangement, together with their proportions and harmonious relations, have been discussed extensively, especially in articles on cosmetic surgery. The description we give below applies equally for human bodies, for which there is also a (different) significant configuration that, again, consists of different elements that are arranged together in precise relationships.

It is generally agreed that faces have a privileged representation in the brain, with at least three areas and possibly more devoted, in part or in whole, to faces. It is also generally agreed that the capacity to recognize a certain “significant configuration” as constituting a face is either inherited or very rapidly acquired, within hours after birth (Goren *et al*., 1975)(Johnson *et al*., 1991), although there has been much discussion as to what it is in the configuration that is instantly recognizable (see discussion in (Zeki & Ishizu, 2013). The preference of the new-born for looking at faces is not found when line drawings of real faces are used (Bushnell *et al*., 1989) emphasizing the pre-eminence of the *biological* concept of face. There are also cortical areas that are critical for the perception of human bodies (de Gelder, 2006)(Peelen & Downing, 2007).

The significant configuration that defines a stimulus as being a face (or a body) constitutes the initial experience and therefore the initial *β prior* from which posteriors are formed, although, unlike colour vision, we have no knowledge of what kind of brain program or algorithm is responsible for determining proportions, harmonies and relationships to specify that the “significant configuration that is being viewed is that of a face. Any individual, regardless of racial or cultural grouping, may assume, with a very high probability, that a significant configuration that one experiences as a face will also be experienced as a face by all members of the human race and therefore that his or her experience has universal validity. Any departures, even minor ones, from the significant configuration that constitutes the biologically inherited or possibly rapidly acquired ^4^ and accepted concept of a face is rejected and never incorporated into the concept of a normal face. Indeed, the cortical response to faces is very exigent in terms of the significant configuration that is accepted as normal and to which there is an optimal cortical response; mis-aligning the two halves of an upright face, for example, delays and increases the typical N170 negative deflection obtained following facial stimulation, as do inverted faces (Ishizu *et al*., 2008); the latter are immediately classified as having an abnormal configuration, and thus not belonging to the biological category of faces. It is also worthwhile to note that the prolonged (1 month) viewing of abnormal or deformed faces does not make them acceptable as normal faces. Indeed, the viewing of stimuli in which inherited concepts of face (and space) are deformed and violated leads to significant activation in fronto-parietal cortex, whereas the viewing of “deformed” or unusual configurations of common objects such as cars or planes do not (Chen & Zeki, 2011). This suggests that such biological concepts are not easily disrupted, a finding that is consistent with our proposed subdivisions of priors into the biological and artifactual categories. Here, it is interesting to note that the painter Francis Bacon, whose self-declared aim was to give a “visual shock”, used deformed faces and bodies to deliver that shock to the spectator; he rarely if ever deformed objects (which fall in to the artifactual category) (Zeki & Ishizu, 2013).

Hence, like with colour categories, an individual can make the reasonable assumption that what they perceive or experience as a face will also be experienced as a face by others and therefore the individual’s experience will have universal validity. Equally, any deformation of a face will be universally experienced as a departure from the significant configuration characterising a normal face. This kind of evidence leads us to suppose that it is very difficult to produce a biologically viable *posterior*, and a belief attached to it, from an inverted face (unlike buildings – or other artefacts – see below) or deformed face; even if produced, it is unlikely to be durable.

### Concept as applied to faces

Our definition of concept as applied to faces is similar to the one given above for colour, *i.e*. an inherited operation or program applied to incoming signals to define them as constituting a face and hence to generate a biological prior from them. Faces and colours share another similarity, namely that they both depend upon ratio taking, though the kind of ratio that is determinant differs between the two. For faces, what must be determined are harmonious relations and constant ratios between its constituents, that is within the face rather than between a face and its surrounds. Hence, we can say that the inherited ratios for faces constitutes a certain significant configuration (Zeki, 2013) in which the various elements (nose, eyes, ears, mouth, eyebrows, *etc*.) bear a constant and pre-determined relationship to one another, a relationship that signifies a face. As the famous Cycladic sculptures testify, it is not necessary that all the elements given above be present at once, but a minimal number of them is mandatory. For example, a minimalist (Cycladic) configuration such as that shown in Figure 2 is sufficient to qualify it as a face. Such a significant configuration then constitutes the inherited initial *β prior* for faces which, as with colour categories, also constitutes the initial experience from which posteriors are formed.

**Figure 2.**
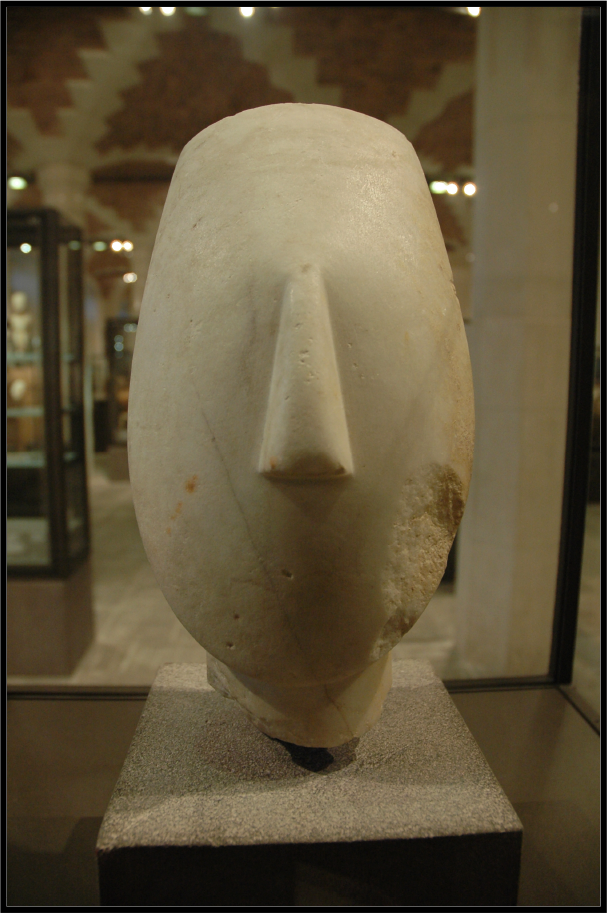
*Head of a female figure, Spedos type, Keros-Syros culture (EC II, 2700– 2300 BC; Louvre Museum, Paris*)

The minimum critical relationships of constituents that determine that a given configuration is that of a face are not known; hence we cannot specify the relationships with the mathematical precision that we can for colour categories. It is nevertheless true that the *posterior* that results from this (face) *β prior* through experience is similarly circumscribed as in colour perception; a significant configuration that constitutes a face cannot be interpreted otherwise, while any departure (as produced by inversions, for example) would mean that the brain will either not classify it as a normal face, or that it will only be temporarily classified as a face, or that it will be classified as an abnormal face, without leaving a permanent posterior.

### The posteriors generated from the *β* prior (significant configuration) for a face

The posteriors generated from a colour category are somewhat confined in scope; for example, those generated from the category green can belong to different shades of green, but do not, in normal circumstances, transmute into another colour category. The posteriors acquired from the *β prior* that constitutes a face are more nuanced. Thus, the posterior can be a face belonging to a particular racial or ethnic group or belong to the category of beautiful or ugly faces (see below). These posteriors depend upon experiencing different faces and therefore upon experimentation. The brain’s Bayesian formula for faces, different from computational facial recognition, is thus similar to that of colours. To signify that one is now dealing with faces instead of colours, what would be needed is to modify the notation from three-dimensional (as in colours) to higher-dimensional case, where the parameter of each dimension would model a specific biological feature, such as the number of eyes, the angle between nose and mouse, the location of the ear, *etc*.

More problematic is whether expressions which convey specific emotions are biologically inherited and hence constitute *β priors*, and whether an individual experiencing a given emotion – for example that of distress – through a particular pattern of facial muscle contractions may reasonably assume that others belonging to the same or a different cultural or ethnic group will experience the same emotion with the same muscular pattern. A facial configuration such as upturned lips, which in fact is universally recognized as a smile, may be significantly dependent upon an inherited template – thus constituting the *β prior* for that expression - and thus not dependent, or only marginally dependent, upon learning and experience. The question may be more complex, however, since the expression of emotion facilitates social bonding and may have a long evolutionary history. Darwin believed that such expressions, involving in each case the interplay of particular groups of facial muscles, are universal and inherited. There has been some controversy in the past over this, but current evidence indicates that there is agreement between different cultures, even those that have not come in contact with each other, about the basic emotions underlying given facial expressions, while there may be some differences regarding the intensity of the emotion conveyed by a given facial expression (Ekman, 1971) (Ekman, 1989)(Ekman *et al*., 1987)(Elfenbein, HA and Ambady, 2002). Evidence suggests that a seven-month old infant has the capacity to recognize facial expressions; given the universality of the recognition, it must be presumed that there is an early interplay here which generates *β priors* according to inherited concepts, possibly augmented by experience and learning, including interacting with family members and others (Nelson and De Haan, 1996)(Safar and Moulson, 2017). This capacity to categorize faces in particular ways – provided they have the significant configuration that constitutes a *β priors* for a face – may thus be compared to the example of colour given above. The colour category generated from a *β prior* comes with an initial hue from which further posteriors are generated; similarly the *β prior* for a face comes when the incoming signals correspond to the brain concept – in the form of a (face) template – that corresponds to a significant configuration signifying a face. To this may be attached an initial particular expression, corresponding to the initial hue in colour vision. The issue here is whether there is a period after birth during which the precise expression is learned, comparable to learning that the hue belonging to a colour category can lead to a new posterior through experience. If so, evidence suggests that that post-natal period is very brief and that the basic emotions conveyed by facial expressions cannot be further modified in their essence by experience and learning.

In summary, conforming to the definition of a *β prior*, an individual who experiences a given significant configuration as that of a face is entitled to assume that others, whether belonging to the same ethnic and cultural grouping or not, will also experience that significant configuration as that of a face. Moreover, an individual experiencing a significant configuration as that of a smiling face or distressed face may also assume that others will also experience that significant configuration in the same way.

### IV a. Face perception and brain physiology

It is generally agreed that there are special areas of the brain that are necessary for the perception of faces, including an area located in the fusiform gyrus known as the fusiform face area (FFA) (Sergent *et al*., 1992)(Kanwisher *et al*., 1997) damage to which leads to the syndrome of prosopagnosia. We note that the FFA is active when faces are viewed from different angles, hence implying a certain degree of *face constancy* (Pourtois *et al*., 2005). Another area critical for faces is located in the inferior occipital cortex, known as the occipital face area (OFA) (Peelen & Downing, 2007)(Pitcher, 2014) while a third area, located in the superior temporal sulcus, appears to be important for the recognition of changing facial expressions (Haxby *et al*., 2000). These may not be the only areas that are important for face perception. It has been argued that the recognition of faces engages a much more widely distributed system (Ishai *et al*., 2005) and that cortical areas classically recognized as face areas may not represent only faces (Haxby *et al*., 2001) since cells responsive to common objects, in addition to faces, can be found in an area such as FFA. Whatever the merits of these contrasting views, they do not much affect our argument, given the heightened susceptibility of faces to distortion and inversion and the relative resistance of objects to similar treatment; this would argue in favour of our general supposition that *β priors* must be separated from α *priors*, whether the representation of objects and faces occurs in the same or in different brain areas (for a general review, Zeki & Ishizu, 2013).

### IV B. The ‘belief’ attached to faces

Hence the biologically based initial belief attached to normal faces must revolve around a configuration that must contain a certain number of features such as eyes, nose, mouth *etc*., set out within certain proportions and harmonious and symmetrical relations to each other which, together, correspond to an inherited or rapidly acquired brain template indicative of faces. There are of course many ways in which faces can be represented; they can, for example, be represented in terms of straight lines in a drawing. But such representations, though recognized as constituting a face, will be immediately classified as a drawing and therefore not a biological face (Bushnell *et al*., 1989). This shows how constrained such a belief and the *β prior* attached to it are. In terms of generality, an individual’s belief that the object one is seeing is a face and that all others will also perceive a face in that configuration is a sound one and makes that belief general. Just like colour, it therefore has universal validity. Extensive experience with distorted faces does not appear to modify the perception of normal faces, thus showing again that *β priors* are resistant to change through experience.

## V. Artifactual (a) priors

By artifactual priors, we refer to the many constructs – from shoes, kitchen utensils, tools of varying complexity to cars and aeroplanes – in brief to man-made products. Unlike colour categories and faces, there is no evidence of a universal, inherited, brain concept, since different cultures have developed different tools and have significantly different architectures. Instead, the brain acquires a concept of these objects through experience and consequent updating of priors, which are strongly culture-dependent. In medieval times, people had no concept of a car or a plane; since their introduction, there have been many modifications of them, and the concepts attached to them have changed accordingly. The concept of a plane that someone living in the 1930s had, for example, did not include jumbo jets equipped with jet engines or variable geometry winged planes; these have been added to the overall concept of a plane since. There are, of course many other examples one could give, including the use of knives and forks and chop-sticks, which differ between cultures and times. Hence one can say that there is no initial, biologically determined prior. Rather, a prior is enabled through post-natal concept formation, which is modified with new experiences.

Crucially, acquired or empirical priors emerge *de novo* and are driven by experiences that are unique to any individual in any given lifetime, although there may be, and usually are, population-level similarities. They therefore are necessarily less precise and more accommodating than biological priors. This follows because they are designed to be modified by experience. The general Bayesian formula for artifactual priors is therefore the classical formulation (see under Bayesian equation) given above, where the priors have a large variance (also see Figure 1).

It is now generally accepted that there is a complex of areas, known as the lateral occipital complex (LOC), which is critical for object recognition (Grill-Spector, 2003). Even though it has been argued that the so-called face areas may not be as specific to faces as originally supposed (see above), and that cells in them may encode objects as well, including ones which we would classify under artifactual categories, the differential response to faces and objects when inverted suggests that they are processed differently (see under faces). Moreover, neural sensitivity to faces increases with age in face-selective but not object-selective areas of the brain, and the perceptual discriminability of faces correlates with neural sensitivity to face identity in face selective regions, whereas it does not correlate with a heightened amplitude in either face or object selective areas (Natu *et al*., 2016).

There is no definitive evidence about when infants begin to recognize objects or whether they recognize faces before recognizing objects. Indeed, it has been shown that infants can recognize differences between shapes even at one month where the outside contour or shape is static and identical, but where the inside smaller shapes are different to each other in each image if, significantly, one of the smaller inner shapes is jiggled or moved (Bushnell *et al*., 1989); this may, in fact, introduce a biological *prior*, that of motion, into the recognition or inference process.

Such results, together with common experience, justify a neurobiological separation between the two categories, faces belonging to the biological category and objects to the artifactual.

### VI. A A biological prior that makes artifactual priors possible?

We have emphasized above the need to separate biological from artifactual *priors* in considering Bayesian operations. Here, it is worth asking whether, at the earliest recorded stages after birth, one can postulate the presence of a *general biological prior* that leads to artifactual priors, which then assume an autonomy of their own (see also Series & Seitz 2013). The common view is that there is one category of cell in the visual brain, the orientation selective (OS) cell (Hubel & Wiesel 1962), which is the physiological ‘building block’ of all forms. This is a plausible argument entertained by both physiologists as well as artists like Piet Mondrian, who defined form as “the plurality of straight lines in rectangular opposition” (see Zeki, 1999). There is an extensive literature on OS cells from birth to adulthood; we do not review this here but summarise this evidence by saying that, while the OS cells and hence the presumed machinery for constructing forms, must be present at birth, they require nourishment in the early stages after birth to mature. Depriving the animal (cat or monkey) at a critical period after birth blights their visual capacities for considerable periods, perhaps even permanently, thereafter. On the other hand, depriving adult animals of vision for equivalent periods in adulthood has little or not effect (Hubel & Wiesel, 1977). Observations in humans deprived of vision at birth through congenital cataracts, with vision restored later in life after successful operations, confirms that visual nourishment during an early ‘critical’ period is necessary for a normal visual life (for a review, see (Zeki, 1993)).

By contrast, a normally nourished visual brain can subsequently recognize and categorize many different shapes or forms, even those that have not been seen before (Logothetis *et al*., 1995)(Freedman *et al*., 2001). Hence, one could consider that OS cells are the given biological *priors*. In accepting the common supposition that OS cells constitute the physiological ‘building blocks’ from which all categories of objects and forms (including faces) are constructed, one must nevertheless acknowledge that (a) OS cells are widely distributed in different, specialized, visual areas of the brain (Zeki, 1978); (b) that the OS cells of V1 may not be the sole source for the neural construction of objects, especially since OS cells in visual areas outside V1 survive the destruction of V1, which suggests that their properties may not be wholly dependent upon input from V1 (Schmid *et al*., 2009); (c) OS cells in different visual areas may contribute to form construction in different ways (Shigihara & Zeki, 2013)(Shigihara & Zeki, 2014a). Moreover, unlike what is commonly posited, V1 is not the sole source of the ‘feed-forward” visual input to the rest of the visual brain; the specialized visual areas, including areas with high concentration of OS cells as well as areas specialized for face and object perception, receive two further “feed-forward” inputs, from the lateral geniculate nucleus and the pulvinar (Cragg, 1969)(Benevento & Rezak, 1976)(Yukie & Iwai, 1981)(Fries, 1981) and are activated with the same latencies, post stimulus presentation, as V1 (Shigihara & Zeki, 2013)(Shigihara & Zeki, 2014b)(Shigihara & Zeki, 2014a). Hence a strictly hierarchical organization for form based on a single feed-forward system from the lateral geniculate nucleus through V1 (as is almost universally supposed), and in which cells within the brain’s form system acquire increasingly more complex properties, enabling them to respond to complex objects and faces, is probably unlikely. Rather, there appears to be multiple hierarchical systems, which operate in parallel and which are task and stimulus dependent (Zeki, 2016). Hence, if the OS cells constitute the biological prior from which forms and objects are constructed in the brain, one must entertain the possibility that there may be several different kinds of OS cells, embedded in different visual areas, each of which may constitute a biological or *β prior* for generating different kinds of posteriors, some of them artifactual. The question has not been addressed experimentally.

There is another difficulty in considering OS as being the sole universal biological *priors*. One cannot build a definitive and exclusive posterior from a single, or from multiple, oriented lines. If faced with either a single or multiple oriented lines, what would the posterior be? This is quite unlike colour, where certain ratios of wavelength composition of light reflected from a patch with given physical properties (reflectance) and from its surrounds determines lead, *ineluctably*, to a given universal biological *prior* in the form of a certain constant colour category, from which certain posteriors, in the form of hues belonging within that colour category, can be elaborated (Land, 1986). We are, we believe, therefore justified in supposing that orientation selectivity, and its manifestation in the OS cells of V1, cannot be the biological *prior* for all forms, as most suppose. Rather, OS cells in different areas may be used to construct different forms or different categories of form, which then act as distinct priors from which posteriors can be generated. But a line need not be a means toward a more complex form; it can exist on its own and in its own right, as artists have so frequently demonstrated. Moreover, there is no belief that can be attached to single oriented lines, except in the narrow sense that they can constitute, either singly or in arbitrary combination, forms in themselves. This is what Alexander Rodchenko (1921) argued when he wrote “I introduced and proclaimed the line as an element of construction and as an independent form in painting”. He added, “…the line can be expressed in its own right, as the design of a hypothetical construction [and can have] a status independent of what is actually taking place, and becomes an abstraction” (Rodchenko, 1921) (our ellipsis). Many artists since then have emphasized the primacy of the line in their work, if not in their writings and the single line or separated single lines are characteristic of the work of many artists, including Alexander Rodschenko, Kazimir Malevich, Olga Rozanova, Barnet Newman, among many others.

Hence, there is no universal belief that is attached to how single oriented lines can be combined. There is also no universal belief attached to what significant configuration of straight lines constitutes a given category of object. The configuration of houses, as places of habitation, differs widely in different cultures – from igloos to huts to skyscrapers and even to inverted pyramidal buildings. One cannot make the assumption that huts are the universal mode of habitation or that inverted buildings depart from the concept of habitation. Rather, the latter are absorbed into the concept (*i.e*. generative models) of habitation through experience (but see below).

### Biological priors in the experience of beauty

Here we sail into more treacherous waters and ask whether there are any *β priors* in one of the most fundamental and universal of human experiences, namely that of beauty. We do so to emphasize the huge range over which *β priors* may operate. We suggest that beauty itself must be divided into biological beauty (in which we include mathematical beauty) and artifactual beauty (see Filippov and Zeki, 2015); it is likely that one will find greater agreement among individuals of different cultures and races within the category of biological beauty (including mathematical beauty – see (Zeki *et al*., 2018)) than within artifactual beauty. The question of beauty imposes itself naturally from one of the main questions addressed in this paper, namely the extent to which individuals can assume that their experience has universal assent. This question is especially important in the context of beauty, widely considered to be a subjective experience, unique to individuals, a belief encapsulated in the old Roman proverb that “*De gustibus et coloribus non est disputandum*” (which incidentally includes color in the subjective category although, as we have argued above, there is little that is subjective in the experience of colour categories given its uniformity). Yet a reasonable inference to be read into Kant’s speculations about the experience of beauty in his *Critique of Judgment* (Kant, 1790) is that any person experiencing beauty in, for example, a face may assume that others will also experience the same face as beautiful. If so, one can also assume that the scope of experience in modifying one’s belief about what constitutes beauty in, for example, a face is also limited though perhaps not quite as limited as for colour categorization.

Contrary to Kant, who believed that some experiences are interfaced through determinate concepts and others through indeterminate ones (see above), we assume that all experiences, including that of beauty, are interfaced through determinate concepts; indeed we do not make a distinction between determinate and indeterminate concepts. But we divide beauty into two categories, though ones that differ from those proposed by Kant. In particular, we posit that the concepts or programs mediating biological beauty are inherited and, just like colour categories, are resistant to change; those mediating artifactual beauty are acquired post-natally, and are therefore hospitable to change. But there is, or there can be, a continuum between the two; this leads us to suggest that some aesthetic experiences may be interfaced through both biological and artifactual concepts, when the experiencing individual can assume universal consent to their experience on a variable scale (see below).

### β priors in the experience of biological beauty

The question that we address here is whether there are any biological priors in the experience of beauty. This is an important, if enormously difficult, subject to tackle. We begin by asking whether there are any universals in beauty that enables an individual from one cultural or ethnic group to assume that what one experiences as beautiful will have universal assent, which is to say that individuals from other ethnic or cultural groups will experience the same thing as beautiful.

It is perhaps best to begin by stating clearly what, in the biological realm, cannot be experienced as beautiful. Faces and bodies are good examples. Starting with what we have stated above, namely that there is a significant configuration which, when it appears in a stimulus, ineluctably leads an individual to identify that object as a face, one can go one step further and say that no face that has two eyes that are placed asymmetrically and in non-harmonious relations with each other (for example, are greatly displaced from one another) can be universally experienced as beautiful (here, we are distinguishing between real human faces as opposed to their depictions in sculpture or painting). Here, an individual can assume with high probability that a human face thus traduced (which implies traducing the inherited brain template for what a face should look like) will not be perceived as beautiful by others, and that what one individual perceives therefore has universal assent.

On the other hand, an individual experiencing a face as very beautiful can also assume with reasonably high probability that their experience has universal agreement. Indeed, it has been shown many times that, broadly, faces regarded as beautiful in one culture are also so regarded as beautiful in other cultures (Langlois *et al*,, 1991)(Langlois Roggman, 1990) (Fink, 2005)(Cunningham *et al*., 1995). Moreover, even infants tend to orient more towards beautiful faces (Langlois *et al*., 1987) regardless of race (Langlois *et al*., 1991). This leads one to suppose that there is a *β prior* for some categories of beauty, in this instance for facial beauty. What is more difficult to determine is what the precise characteristics determining facial beauty are, beyond the generally accepted proposition that they involve symmetrical disposition and harmonious relations of constituent parts. It is perhaps easier to define what does not constitute facial beauty; in general terms, it involves the subversion of the inherited template for a significant configuration for faces – a fact well recognized by the painter Francis Bacon (see above).

Hence, we propose that there is a basic significant configuration which constitutes the initial *β prior* for a face and, at the same time, the initial *β prior* for beauty on a face, the precise characteristic of the latter being ill-defined but universally recognized in terms of the experience of beauty. The parallel to this in colour vision is the *β prior* that determines both the colour category and the initial hue attached to that category. From these biological priors posteriors can be generated through experience, as indeed they seem to be over months post-natally, when infants become more familiar with faces (Lee *et al*., 2017). But, what confers on a normal face the inestimable quality of beauty is much more difficult to define. In his *Kanon*, the Greek sculptor Polykleitos suggested that the perfect human body (such as his *Dorypherous*) can be sculpted by using a body part such as a phalanx and treating it as one side of a square; when rotated the square’s diagonal gives a ratio of 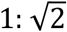 which can be applied, by multiplication, for creating the next phalanx and then repeating the procedure until the whole forearm is sculpted; in the final product, the different and contiguous parts bear a definitive mathematical relationship to each other. While agreeing, in his *Vier Bucher von menschlicher Proportion*, that a canon of mathematical proportions must be applied to the geometrically constructed human face and figure, Albrecht Dürer believed somewhat more realistically that the beauty of form was relative and not absolute. Similarly, Leon Battista Alberti believed in averages or means when constructing the perfect human figure, as when he states in *De Statua* that, “I proceeded accordingly to measure and record in writing, not simply the beauty found in this or that body, but, as far as possible, the perfect beauty distributed in Nature, as it were in fixed proportions, among many bodies” (Aiken, 1980), a supposition that finds some support in infants’ preference for the averageness in faces (Aharon *et al*., 2001)(Loffler *et al*., 2005)(Anzures *et al*., 2009). Hence, while proportion and harmony are generally agreed to be pillars of beauty in the human figure and face, there is no agreement or even knowledge of absolute values but rather a tendency towards average or mean values. Whether there are absolute or mean values, the general point we are making is that, for a face to be immediately recognized, first as a face and next as a normal face and finally as a beautiful face (see also below), not only must at least some of its constituents be present but they must be there in certain, fairly precisely defined, mathematical relationship to one another.

### The experience of artifactual beauty

There are many examples of artifactual beauty, which include all man made objects; such objects are interfaced through brain concepts which are acquired post-natally and continually updated, thus making them more hospital to Bayesian analyses. It is likely, for example, that those brought up in Oriental cultures will find greater beauty in the noble temples of the East and those nourished in Western culture in the great cathedrals and abbeys of the West. Yet, when it coms to artifactual beauty, it is also interesting to question whether the experience of artifacts such as buildings are not also interfaced through biological concepts, in addition to acquired, artifactual ones. This can be illustrated by reference to architecture and architectural beauty in general. It would be surprising if the newborn has a concept of a house or some building that serves a given purpose. The concept of a building is acquired post-natally as are concepts of different kinds of buildings, for example a habitation, an industrial building or a temple or church. The question becomes more difficult when one enquires whether there is any determining concept regarding beauty in architecture, which amounts to asking whether architectural beauty belongs to the biological or artifactual category, or to both. The answer may not be quite straightforward.

### The categorization of architectural beauty

Although we have included in the artifactual category human artefacts such as cars, utensils, and buildings, the separation between biological and artifactual priors is in fact more nuanced. We can illustrate this by asking whether there is any determining concept regarding beauty in architecture, which amounts to asking whether architectural beauty belongs to the biological or artifactual category. An ancillary question is to ask to what extent an individual belonging to a certain ethnic or cultural group can reasonably assume that what one considers to be a beautiful architectural design will have universal assent, in the sense that others will also experience that architectural design as beautiful?

The architect as well as the viewer of buildings would have already looked at nature and something of that (biological) experience would have surely seeped into the experience of both. The Roman architect Vitruvius (for whom beauty or *Venustas* constituted one of the three pillars or Triads of architecture) emphasized that beauty in architecture is derived at least in part from contemplating the beauty of nature, and especially the beauty of the human body the experience of which, just like the experience of human faces, is interfaced through inherited brain concepts (see above). Like Vitruvius, Leonardo Da Vinci also believed strongly in copying nature when trying to represent objects and their beauty. In his *Vitruvian Man (Canon of Proportions)* he worked in the reverse direction from Vitruvius: he adapted mathematical architectural principles, derived from the Vitruvian architectural principles, to produce the perfect human body, which represented for him, in miniature form, the harmonies and proportions present in nature and the Universe, and which he considered to be a “*cosmografia del minor mondo*”. Hence, if architectural principles, derived from inherited universal laws of deductive logic that are at the basis of mathematics (Zeki *et al*., 2014)(Zeki *et al*., 2018), are used (consciously or unconsciously) in the creation of perfect human bodies for an artist like Leonardo, these same principles can also act in the reverse direction too – in influencing architectural design. Indeed, one is tempted to read into the principle of *pareidolia* the desire of the architect to instil, unconsciously, properties derived from more biological percepts such as those of faces or bodies or landscapes into architectural design. It is indeed common to find many architectural designs that are inspired by, and resemble, human bodies or body parts.

It is therefore reasonable to suppose that there is a heavy dose of biological beauty, dependent upon inherited brain concepts, that regulates architectural design, provided it is not subject to other requirements, as detailed above. We therefore imagine, though we cannot be sure, that, if forced to do the experiment, our outside observer would find that, even though the unanimity is not nearly as great as those found in colour categorization, there is greater unanimity in classifying buildings as beautiful than is commonly supposed; hence that, even in the domain of architecture, beauty is not quite as subjective as may seem at first. Although there are many considerations that go into architectural design, what universality architectural beauty may possess probably lies in satisfying inherited brain concepts of proportion, harmony and geometric relationships that are more formally expressed in mathematical beauty.

## VII. Conclusion

We have here given a general account of what we believe is an important distinction to be made when considering the brain as a Bayesian system. For simplicity, we have concentrated on extreme examples, ones which we have better knowledge of, namely that of colours and faces for the *β priors* and of common artefacts such as houses for the *α priors*. This naturally leaves out of account a vast territory in which both priors may be involved. Laplace himself delved into questions of average mortality and the average duration of marriages. The list can be extended to include social interactions as well as economic activity in which the unfortunately un-studied *β prior* of greed may play a crucial role, in addition to *α priors*. An example of the latter, which plays a role in economic calculations, is the recognition of political decisions that influence monetary values, which would fall into the artifactual, α, category. In these, and many other human activities that involve making inferences based on a set of beliefs, the distinction between the two categories of priors is, we believe, important. Finally, the distinction between biological and artifactual priors can also be extended to aesthetics, since aesthetics pertaining to biological entities such as faces or bodies, are similarly constrained by the configurations that constitute them (Zeki, 2009)(Zeki, 2013).

We have restricted ourselves here largely to the visual brain, but hope to deal with other brain processes that are subject to Bayesian-Laplacian operations in future papers.

## Acknowledgement

We are very grateful to Karl Friston, Will Penny, Stewart Shipp, and Tianchen Qian for their insightful comments on an earlier version of this paper.

## Supplementary Information

The central point we are making in this article is that there exist two categories of priors, artifactual and biological, the latter of which is resistant to modification by experience (or experimentation). Below, we supplement our arguments given above in the language of Bayes theorem.

Accommodating the definition outlined in Figure 1 of the text, let us consider a quantity of interest μ. In color vision, this quantity consists of three items of information (namely the long, medium, and short wavelengths, respectively); hence μ is a three-dimensional vector. For face and object recognition, the dimension is much higher (because there are more parameters that define a face or an object), although the underline arguments are similar. We proceed, without loss of generality, presuming μ is a vector in 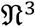.

For simplicity, assume *μ* follows a multivariate Gaussian distribution with a mean and covariance *μ*_0_ and *∑*_0_, namely, *μ* ~ *MVN*(*μ*_0_, ∑_0_). Suppose we conduct *n* experiments (of viewing a patch), and obtain *n* sets of data (*x*_1_, *x*_2_,…,*x*_*n*_), each being a three dimensional vector containing the ratios of long, medium, and short wavelengths reflected from the centre and the surrounds in each experiment *i*. The observed data also has a distribution (for simplicity, we assume it is a multivariate Gaussian distribution with mean and covariance *μ* and *∑*, namely, *x*_*i*_ ~ *MVN*(*μ*, *∑*). It follows that, algebraically, the posterior distribution for *μ* is

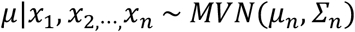

where 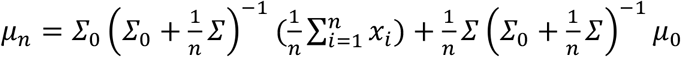 and 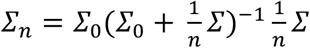.

If a prior is biological, then it means that the variability, or *∑*_0_, is very small, namely very close to **0** (where **0** indicates a three by three matrix with all zero entries), and the posterior mainly dependent upon the prior and independent of the observed data (*x*_1_, *x*_2_,…,*x*_n_), in the sense that the posterior mean *μ*_n_ is very close to the prior mean *μ*_0_, and the posterior covariance *∑*_*n*_ is very close to ***0*** (in other words the sum of all absolute entries of *∑*_*n*_ is close to 0). In the extreme case when *∑*_0_ = ***0***, we have the posterior mean being identical to the prior mean (*μ*_*n*_ = *μ*_0_), and posterior covariance *∑*_*n*_ = ***0***; in other words, when there is no variability in the prior, the posterior is solely dependent on the prior and not at all on the observed data (or experimentation) (see Figure SI (a)-(b)).

When a prior is artifactual, where it has a variability that is reasonably larger than zero, the posterior takes information from both the prior and the observed data. More specifically, using the Sherman–Morrison–Woodbury formula and some matrix algebra, we can further write

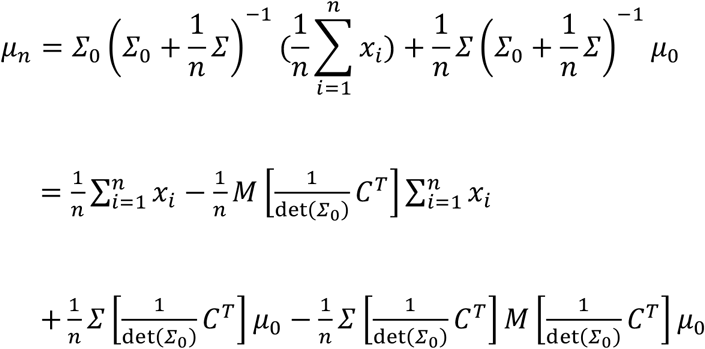

and

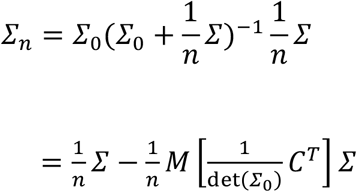

where 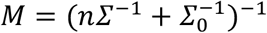 and *C* denotes the *adjugate* of *∑*_0_.

Critically, when the variability of the prior *∑*_0_ is large (in the sense that the determinants det (*∑*_0_) is large), the posterior mean *μ*_n_ is dominated by the observed data (namely 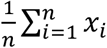; particularly, when det 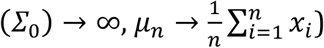 and the posterior covariance is dominated by the covariance of the observed data (namely 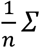; particularly, when det 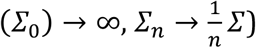. Put in a different way, when we do not have much information from the prior (indicated by its large variability), the posterior tends to learn information solely from the observed data.

The above mathematical arguments are illustrated in the figures below.

---

**Figure SI:**
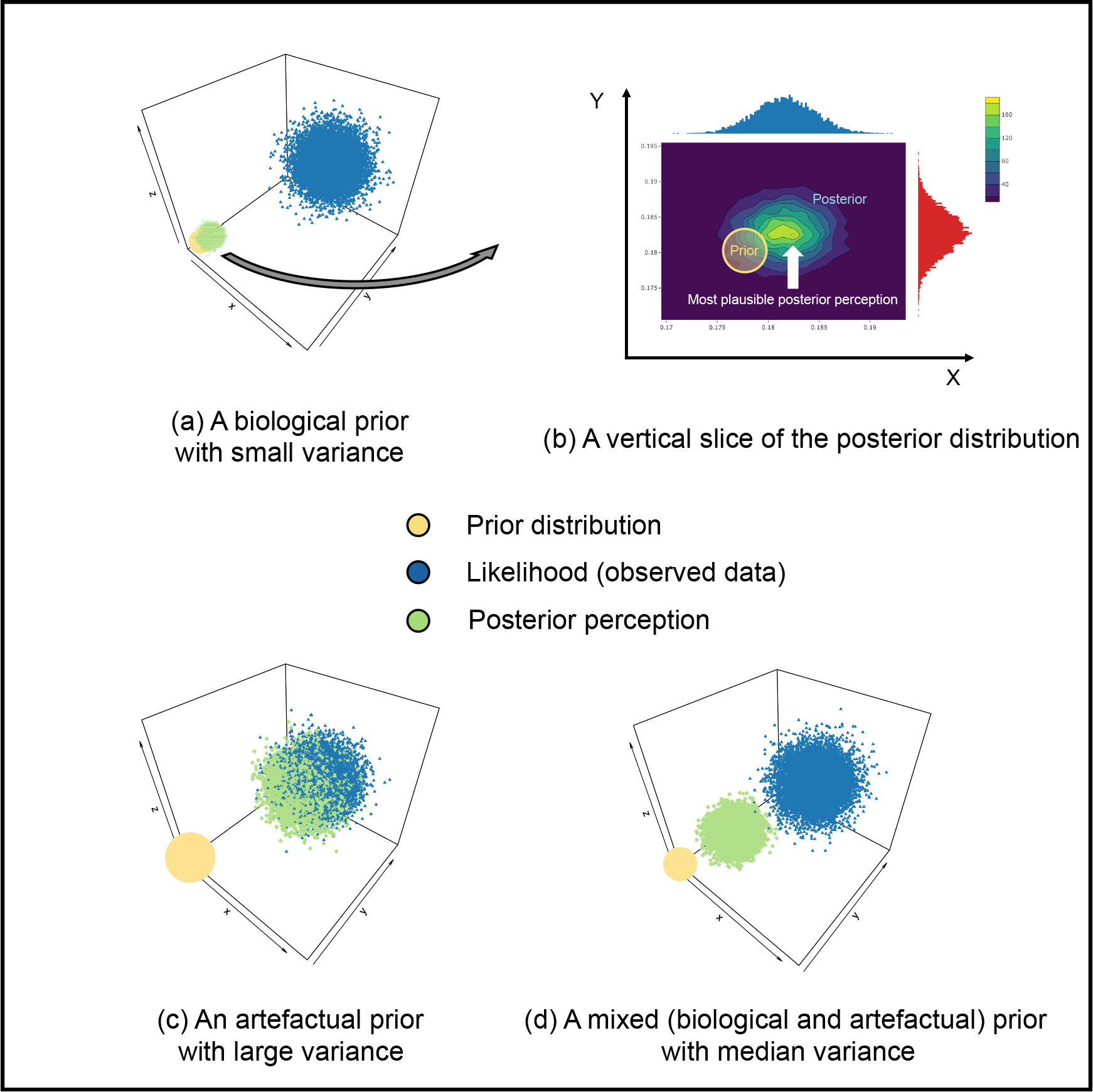
An illustration of the relative resistance of biological priors to modification through experience compared to artifactual priors. Priors are represented in yellow, observed data (experience) in blue, and the posterior produced from observation in green. Size of circles indicates the extent of variability. For comparison purposes, the observed data (blue dots) in (a), (c) and (d) are simulated using the same (multivariate Gaussian) distribution. (a) When the prior distribution (indicated by the yellow dots (see figure (b) for a zoomed-in snapshot) has very small variability, the posterior distribution is derived mainly from the prior, and not much from the observed data. This constitutes the concept of a biological prior. (b) shows that, with biological priors, the centre of the posterior distribution is very close to that of the prior distribution. (c) shows that when the prior distribution (yellow dots) has a very large variability, the posterior distribution (indicated by the green squares) is modified mainly by the observed data (blue dots). This constitutes the concept of an artifactual prior. Note that the centre of the posterior distribution in (c) is very close to the centre of the observed data (blue), under an artefactual prior. (d) When the prior distribution (yellow) has moderate variability, the posterior distribution is derived from both the prior and the observed data; the centre of the posterior distribution is between the prior distribution (yellow circle) and the distribution of the data (blue).

## Footnotes

3 For brevity, we restrict ourselves to long, middle and short-wave light, without giving the peak values along the spectrum; in practice, and under natural viewing conditions, a surface will reflect light of many wavelengths, but there will be a (constant) ratio for light of any wavelength reflected from the centre and surrounds.

4 Note that the term “rapidly acquired” also signifies some inherited element, since the acquisition spoken of here enjoys a privilege over other concepts, such as those of buildings, which are not as rapidly acquired.

